# The outer membrane proteins OmpA, FhuA, OmpF, EstA, BtuB and OmpX have unique lipopolysaccharide fingerprints

**DOI:** 10.1101/448274

**Authors:** Jonathan Shearer, Damien Jefferies, Syma Khalid

## Abstract

The outer membrane of Gram-negative bacteria has a highly complex asymmetrical architecture, containing a mixture of phospholipids in the inner leaflet and in the outer leaflet they contain almost exclusively lipopolysaccharide (LPS) molecules. In *E. coli,* the outer membrane contains a wide range proteins with a beta barrel architecture, that vary in size from the smallest having eight strands to larger barrels composed of twenty-two strands. Here we report coarse-grain molecular dynamics simulations of six proteins from the *E. coli* outer membrane OmpA, OmpX, BtuB, FhuA, OmpF and EstA in a range of membrane environments, which are representative of the *in vivo* for different strains of *E. coli.* We show that each protein has a unique pattern of interaction with the surrounding membrane, which is influenced by the composition of the protein, the level of LPS in the outer leaflet and the differing mobilities of the lipids in the two leaflets of the membrane. Overall we present analyses from over 200 microseconds of simulation for each protein.

**Author summary:** We present data from over 200 microseconds of coarse-grain simulations that show the complexities of protein-lipid interactions within the outer membranes of Gram-negative bacteria. We show that the slow movement of lipolysaccharide molecules necessitate simulations of over 30 microsecond duration to achieve converged properties such as protein tilt angle. Each of the six proteins studied here shows a unique pattern of interactions with the outer membrane and thus constitute a ‘fingerprint’ or ‘signature’.

## Introduction

The outer membrane proteins (Omps) of Gram-negative bacteria perform a wide range of functions including signalling, host cell adhesion, catalysis of crucial reactions, and transport (active and passive) of solutes/nutrients into and out of the cell. Given the range of functions they perform, it is not surprising that Omps have a wide range of sizes and oligomeric states. Currently the structures of > 70 Omps from Gram-negative bacteria have been determined; all of these with the exception of, Wza from *E. coli,* have beta barrel architectures (see http://blanco.biomol.uci.edu/mpstruc). In *E. coli,* the sizes of Omps range from barrels composed of eight beta strands to twenty-two strands for a monomer[1, 2]. The structure-function relationships of a number of Omps have been the focus of experimental and simulation studies over the last twenty years or so, and consequently many details have emerged, some of these are reviewed in [3, 4]. More recently it has become evident that the interactions of Omps with lipids within their local membrane environment can be of a specific nature, these interactions are therefore likely to have functional consequences. For example, experimental studies have shown that OmpF has a specific LPS-binding region on its outer surface. In addition to their interaction with lipids, Omps also interact with each other as well as with peripheral membrane-binding proteins, which is perhaps not surprising given the crowded nature of the outer membrane. However, as with lipid-Omp interactions, Omp-protein interactions are also often specific and have functional consequences that may impact upon the life-cycle of the bacterium. For example, it is known that the translocation of vitamin B12 across the outer membrane is facilitated by binding to BtuB, and then the subsequent formation of a translocon complex with OmpF. To understand these interactions and how they are stabilised, it is important to characterise the orientation and dynamics of Omps in their native membrane environment, as well as the impact they have on the local membrane.

Molecular dynamics simulations at fine-grain and coarse-grain levels of resolution have been employed to study the orientations of many membrane proteins. The most comprehensive study is perhaps that of Stansfeld *et al* in which coarse-grain simulations of membrane proteins embedded in symmetric phospholipid bilayers are performed and the results deposited in an online database[5]. The study covers all known structures of membrane protein and is updated as more structures become available. For Omps, however the analyses is somewhat complicated by the biochemistry of the outer membrane; the two leaflets differ in their lipid composition. The outer leaflet contains lipopolysaccharide (LPS) and the inner leaflet contains a mixture of zwitterionic and anionic phospholipids. Given that the length of the LPS molecule can vary, and it differs from phospholipids, it is important to ascertain if this impacts upon the orientation of Omps. We and others have previously shown that inclusion of this biochemical heterogeneity is important for understanding the conformational dynamics of Omps[6-9]. However a thorough study of the interplay between these proteins and their local membrane environment and how this may differ in a protein-dependent manner, such as the study by Tieleman and co-workers in which the ‘fingerprint’ of each protein is identified, has so far not been reported for Omps in their natural lipid environment[10]. One particular technical issue that arises when simulating LPS-containing membranes is that of conversion, given the slowly diffusing nature of the lipids. It has been shown previously that atomistic LPS-containing membranes require simulation on the order of 500 ns for equilibration[11].

To identify whether the patterns of Omp orientations, and their interactions with the local membrane environment are uniform or dependent upon the protein and/or membrane type, we have performed coarse-grain simulations of six Omps of varying sizes, namely OmpX, OmpA (N-terminal domain), BtuB, FhuA, OmpF and EstA in outer membrane models that vary in the level of LPS in the outer leaflet. This set of proteins has been chosen as it includes small (OmpX and OmpA), large, (BtuB and FhuA) multomeric (OmpF) and multidomain (EstA) Omps. We find that the level of LPS, but also the differences in mobility of the two leaflets of the outer membrane impact the protein-membrane interactions of the six Omps in different ways. Thus to use the terminology of Tieleman and co-workers, each one of these proteins has a unique LPS ‘fingerprint’. In undertaking this study we have also been able to determine the timescales that are required for convergence of various properties of LPS-containing systems; these are significantly longer than those needed for simpler phospholipid membranes.

## Results

For clarity, the results are divided into two broad categories; Omps in varying levels of LPS and Specific-Omp interactions, each of which is subdivided into smaller sections. The four different membrane environments have the same inner leaflet composition (90% 16:0–18:1 phosphoethanolamine (POPE), 5% 16:0–18:1 phosphatidylglycerol (POPG) and 5% cardiolipin), whereas the outer leaflet is composed of one of the following: either 100 % rough lipopolysaccharide at (i) Re level, termed ReLPS (ii) or Ra level, termed RaLPS (iii) or smooth LPS which incorporates 5 sugar units of O-antigen, termed OANT (iv) or a mixture of smooth LPS and POPE in a 4:1 ratio, termed OANT_PE, further details are given in the methods section.

Table 1 provides a summary of the simulations that have been performed for this study. For each of the six proteins, at least 2 × 30 microseconds simulations have been performed of the protein in 4 different environments. Giving a simulation time of 240 microseconds per protein. Inevitably with this type of study, see for example Stansfeld et al, Scott et al, Corradi et al, an enormous amount of data is generated, and it is not possible to present or discuss comparative analyses for all proteins across all metrics[5, 10, 12]. Therefore we have presented the analyses for all protein in the supplementary information and have compared interesting cases in the main text of the paper.

**Table 1.**
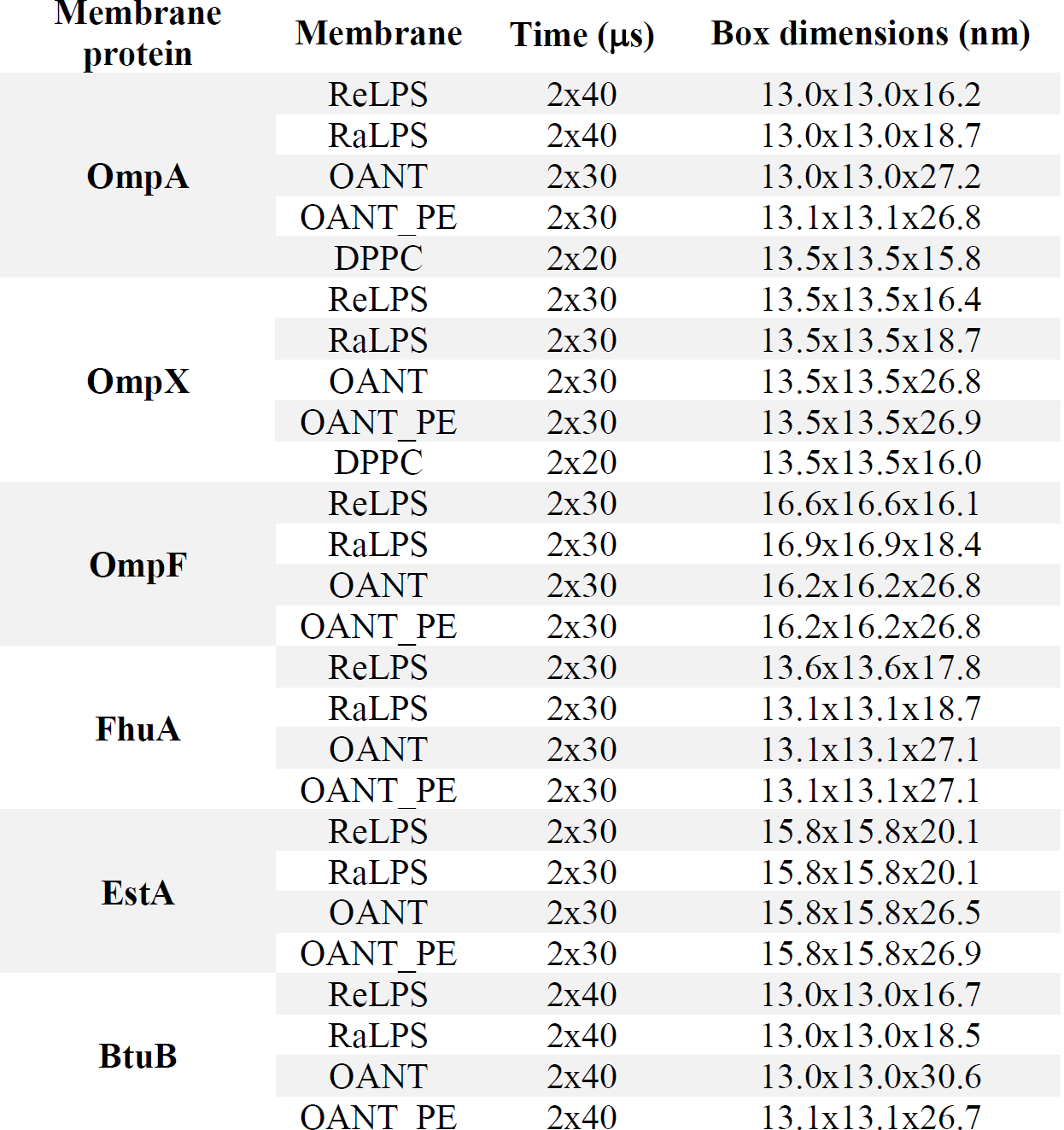
Details for all of the simulations carried out for this paper. The box dimensions refer to the initial size of each system generated by CHARMM-GUI.

### Orientation of Omps in varying levels of LPS

Molecular Dynamics is an established technique for prediction of protein orientation within membranes. In particular the smoother energy landscapes and longer timescales accessible to coarse-methods make them especially useful for this purpose[13]. As has been demonstrated previously, for symmetrical bilayers they can be used to self-assemble a bilayer around the protein, thus removing any bias in terms of human intervention in determining the orientation[12]. For asymmetric bilayers, self-assembly is not straightforward as one cannot control the composition of each leaflet. But the faster dynamics afforded by coarse-grain models do provide a route to determining equilibrium protein orientation that is independent of the initial orientation[5]. We have calculated the tilt angle with respect to the membrane plane of all six proteins in the four model membranes summarised in Table 1 (Figures 1 and S1).

**Figure 1.**
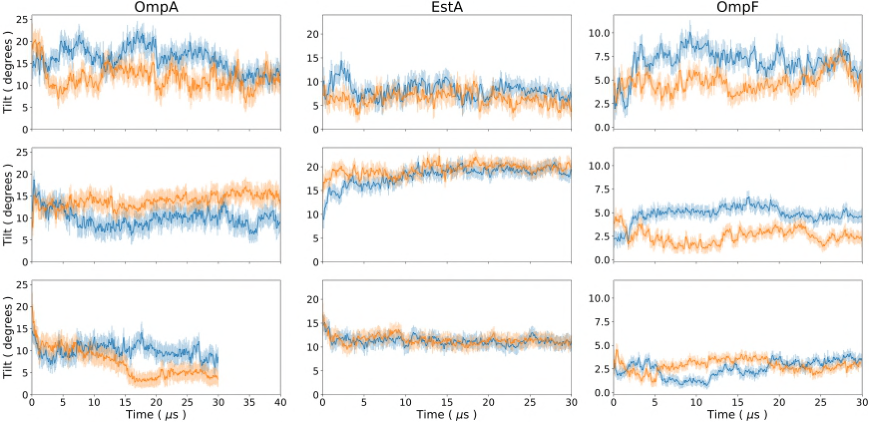
The protein tilt of the transmembrane region of each protein backbone with respect to the z axis as a function of time. In the case of OmpF the tilt was determined using the entire protein backbone. Each column is labelled with the protein that all tilts in that column correspond to. Each row of the above plot corresponds to a certain level of LPS: ReLPS, RaLPS, OANT are in descending order. Each line in a subplot corresponds to a single repeat. The errors were generated from the standard deviation of block averages taken over 100 ns intervals.

Across two independent repeats of each simulation, convergence is largely achieved after 30-40 μs, with the O-antigen containing membranes showing the greatest variation between independent repeat simulations. Our results indicate that the proteins differ in the way they are orientated depending on the level of LPS in the membrane. Furthermore, compared to simple phospholipid membranes, the Omps can access a wider range of angles in LPS-containing membranes. Interestingly, the magnitude of differences in tilt angles, and pattern of tilt versus LPS type varies from protein to protein. For example, if we consider the two eight stranded barrels, OmpA and OmpX (Figures 1 and S1B), then we note that OmpA is tilted at an angle of ~12 °(where 0 ° and 90 ° indicate the protein is perpendicular and parallel to the plane of the membrane respectively), in ReLPS and O-antigen level of LPS. In RaLPS there is less overlap of the two repeat simulations, but the trends indicate a tilt of ~10-13 °. Interestingly in the OANT_PE system OmpA is tilted at an angle of ~8 °, so there is no simple correlation between tilt angle and length of the LPS molecule. OmpX, is tilted at about 5-7 ° in ReLPS, ~12-13 ° in RaLPS, 8-9 ° in OANT and in OANT_PE the two independent simulations vary without overlapping. Similarly for the other four proteins, there is no simple correlation between LPS length and protein tilt. The protein EstA is the only Omp in our test set that has an extracellular domain large enough to interact with all sugar sections in each LPS type. The EstA tilt angles are about 5, 20 and 10 degrees in ReLPS, RaLPS and OANT systems, respectively (Figure 1). The increased tilting in RaLPS systems seems to be the result of the extracellular bulb-like domain tilting to form interactions with the core oligosaccharides. In ReLPS systems the bulk domain is too far from the sugars for tilting to occur; whereas in O-Antigen systems the level of tilting observed in RaLPS is unnecessary (Figure S2) as stabilising interactions form with the O-Antigen chains. Thus, from our results it seems that for Omps with substantial extracellular domains, the level of LPS can play an important role in the membrane alignment of the protein.

Here it is important to consider an aspect of the outer membrane asymmetry that is usually neglected in the context of Omp orientation; mobility of the lipids within the two leaflets. The LPS molecules in the outer leaflet are less mobile than the phospholipids of the inner leaflet; this is now well-established[4, 14]. To understand how this impacts on the motion of smaller Omps in each leaflet, we considered the proteins OmpX and OmpA. We divided the barrels of each protein into the region that is contained within the outer leaflet and the region that is found in the inner leaflet. We calculated the centre of geometry motion of each portion of the barrel for time period 0-30 ns (Figures 2 and S3). It can be clearly seen that in general, the portion of the protein in the inner leaflet samples more of the membrane than the portion within the outer leaflet, but the trend is more marked in the smaller proteins. If we consider OmpX in detail (Figure 2); during the first 5 microseconds of simulation there is greater movement of the protein within the inner leaflet compared to the outer leaflet, while during the last 5 microseconds as an equilibrium orientation is approached, the amount of movement within the two leaflets is comparable. For reference we calculated the centre of geometry motion of OmpX and OmpA in simple phospholipid bilayers (Figure S4); where the differences in the amount of the membrane traversed by the protein in each leaflet is far smaller. Taken together the results of the tilt and motion analyses indicate that although it has been suggested for many years in the literature that Omp location and orientation within the outer membrane are governed by aromatic residues on their outer surfaces which are positioned at the lipid headgroup/tail interface, matching the hydrophobic surface of the Omp with the hydrophobic region of the surrounding bilayer respectively, certainly the latter is rather an oversimplification for the Omps with smaller barrels.

**Figure 2.**
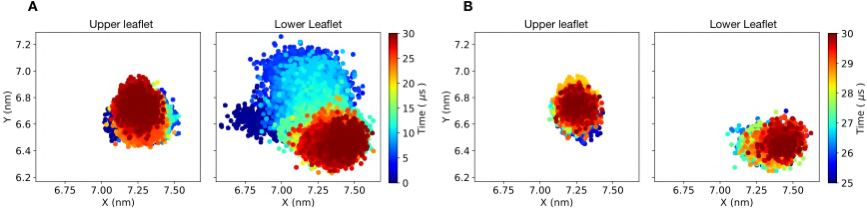
The barrel of OmpA was divided into regions that were in either the lower (see right) or upper (see left) leaflet of the ReLPS outer membrane based on the center of geometry of the entire transmembrane region of the protein backbone. The center of geometry of the protein backbone of each region was then plotted as a function of time for A) 0-30 ps and B) 25-30 ps. Before this analysis was carried out each trajectory was centered on the protein.

### Specific Omp-lipid interactions

The hydrophobic regions of all our model outer membranes have the same width, as the lipid A component is unchanged, therefore given we see differences in their behaviour in the different membranes, interactions with the sugar moieties of LPS clearly have an influence on the protein orientation and location. To probe specific protein residues that form stabilising interactions with the lipids of the outer membrane, we have calculated the number of interactions of all protein residues with the LPS headgroups and sugars. Give the headgroup of LPS is the same in all membrane models, we analysed only the ReLPS containing OM model for these interactions. In general we observe that aromatic and basic residues have greatest propensity to interact with the LPS headgroups (Figures 3 and S5A). The interactions with aromatic residues are in agreement with many previous studies of Omps in phospholipid bilayers, and are posited to anchor the proteins in the outer membrane. In LPS we observe these interactions to be dominated by tyrosine residues, although in OmpA the numbers of interactions with tyrosine and tryptophan are comparable. The headgroup interactions with basic residues are largely with the phosphate particles of the LPS headgroups (Figure S5B) as would be expected on the basis of electrostatics. Each protein has its own distinct pattern of interactions; in OmpX and OmpF interactions of the basic residues with LPS headgroups are largely though Lysine residues whereas for BtuB and FhuA the number of interactions are comparable between arginines and lysines and for EstA there are more interactions with arginine than lysine. Intriguingly for OmpA, interactions with asparagine and histidine dominate. The extracellular loops of OmpA and OmpX both form contacts with the sugars and the lipid A headgroups (Figure S6). Closer inspection of the composition of the extracellular loops in OmpA reveals a reduced number of positively charged amino acids compared to OmpX. Previous atomistic studies have reported that loops rich in positively charged amino acids in OprH from *P. aeruginosa* mediate interactions to the lipid A headgroup[7]. OmpA dimers have recently been characterised, although their biological significance is still debated. We note here that the loops of OmpA are known to define the interface of the OmpA dimer through polar and non-polar interactions, therefore having basic residues in these loops would likely alter the propensity of the protein to dimerise/destabilise the dimerization interface[15, 16]. The dimer was not studied here as OmpA is thought to be present mostly in the monomeric form.

**Figure 3.**
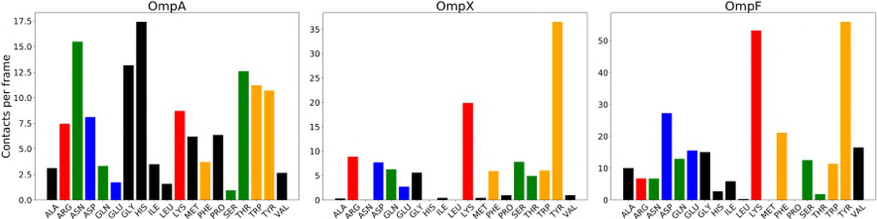
Average number of contacts per frame between each residue in a protein and the Lipid A headgroup for 25-30 ps across both repeats. Each bar was coloured with respect to the polarity of each residue: red=positively charged, blue=negatively charged, orange=aromatic, green=polar and black=non-polar.

We next sought to quantify the interactions of the six proteins with the sugars of LPS (Figures 4 and S7). Once again these interactions are largely dominated by basic residues in ReLPS, with the exception of BtuB, however they are more variable in RaLPS and O-antigen. For example if we compare BtuB and FhuA given their size and functional similarities we may expect their interactions with LPS to be consistent with each other. We find in BtuB the highest frequency of contacts are with tyrosine residues in all membrane types. The highest number of contacts with charges residues are with lysines in ReLPS, aspartates in RaLPS and comparable between lysines and aspartates in OANT. On the other hand, in FhuA the highest frequency of contacts at the ReLPS level are with histidines closely followed by lysines whereas in RaLPS and OANT contacts with lysines dominate. The interactions with sugars of the multidomain protein, EstA, also show variations in the polarity of the charged interactions (Figure 4); they are largely through arginine and aspartate residues in ReLPS, but with lysine in RaLPS and then again with aspartate in OANT. This can be explained by considering the distribution of residues. There is a cluster of five lysine resides in the ~ middle section of the extracellular domain (Figure 4). This cluster of lysines can form contacts with the sugars in RaLPS, but is not accessible to ReLPS. Likewise many of the aspartate residues that form interactions with sugars in the OANT membrane are not accessible to RaLPS. Thus, the pattern of contacts with charged residues observed for EstA can be explained by a comparative deficiency of lysine in the upper and lower regions of the extracellular domain of EstA. Overall, the protein-LPS contact data show that the pattern of contacts is not general (for the six proteins we have studied here), the interactions may be dominated by charged residues which can be acidic or basic, or they can be dominated by polar or aromatic residues, precisely which, is dependent on the Omp but also the level of LPS present in the outer leaflet of the outer membrane. As a validation of our methods it is important to compare to experimental data, where this is available. Experimental details regarding the interactions of FhuA and OmpF are available in the literature; one of the X-ray structures of FhuA is resolved with a lipid A molecule bound to the protein[17] and OmpF has also been shown to have specific LPS binding sites[18]. Our simulations identify the same region of FhuA as making contacts with LPS, as that bound to Lipid A in the X-ray structure (Figure 5). Lakey and co-workers have shown that OmpF requires LPS to form a timer *in vivo* through the binding of LPS to positively charged amino acids[18]. Our simulations are consistent with these observations, with OmpF interactions with LPS sugars being dominated by lysine residues (Figure 4). The interactions with LPS headgroups are dominated by lysine and tyrosine residues (Figure 3). This is consistent with the X-ray structure of the porin OmpE36 in complex LPS, determined by the same authors which shows interactions between LPS headgroups and Lysine and tyrosine residues [18].

**Figure 4.**
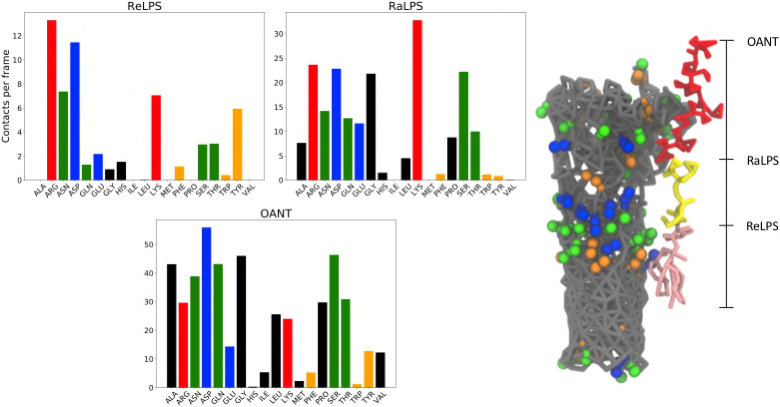
(left) Average number of contacts per frame between each residue in EstA and the lipopolysaccharide (LPS) core oligosaccharides and O-Antigen (if present) from 25-30 ps across both repeats. Each row corresponds to a given protein, while each column corresponds to the LPS system the protein was embedded in (see Table 1). Every bar was coloured with respect to the polarity of each residue: red=positively charged, blue=negatively charged, orange=aromatic, green=polar and black=non-polar. Note that multiple contacts between a single residue and another Lipid A headgroup were only counted once. (right) Model of EstA backbone (grey) with all Arginine (orange), Aspartate (green) and Lysine (blue) residues shown next to a smooth LPS lipid. The height of LPS in ReLPS (pink), RaLPS (yellow) and OANT (red) systems is marked next to the LPS model.

**Figure 5.**
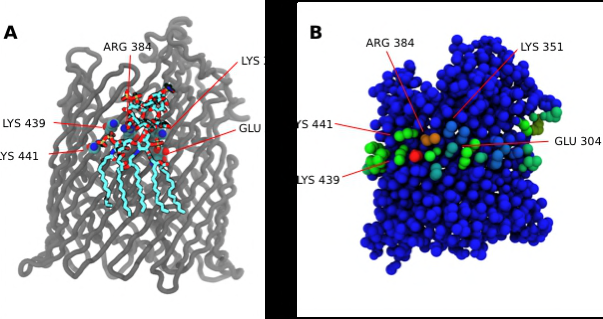
A) Image of crystal structure of FhuA (pdb code = 1QFG) which contains a bound lipopolysaccharide (LPS). The marked residues where within 0.4 nm of the Lipid A phosphate groups. B) Contact map of the coarse-grained (CG) structure of FhuA, between each FhuA residue and the phosphate beads of Lipid A in a ReLPS membrane. The contact analysis was carried out from 25-30 ps using a cutoff of 0.6 nm (as is standard for CG models). Each residue was coloured based on their respective number of contacts from low (blue) to high (red). The residues identified in Figure 5A are marked for the sake of comparison.

Thus far the analyses have been protein-focussed; but it is just as important from the perspective of the structure-dynamics-function relationship, to ask how the membrane responds to the proteins. To this effect we have analysed the enrichment/depletion of lipids around each Omp, the membrane thickness, and the effect of the proteins on the orientation of the O-antigen polysaccharide chains.

### Membrane thickness and lipid enrichment/depletion

Previous simulation studies have shown that Omps have a tendency to cause local thinning of phospholipid bilayers[12, 19]. Given the lipid A tails are shorter than the tails of the phospholipids used in both leaflets of the bilayer in previous studies, it is of interest to determine whether Omps induce also induce thinning of the more biologically relevant models of the outer membrane used here. Indeed, we find the membrane to be thinner within ~2.5 nm of all six proteins, in all 4 outer membrane models, beyond which can be considered ‘bulk’ membrane (Figure S8). This is the same distance from the protein that membrane thinning and thickening effects were reported for a range of beta barrel and alpha helical membrane proteins by Scott *et al* in phospholipid bilayers and for helical membrane proteins by Corradi et al, in our case we find the effect is always thinning of the membrane around the protein, regardless of the level of LPS. It is of functional relevance to explore the lipidic origins of the thinning effect; is it a general feature of membranes around Omps, or a function of specific lipid types? In order to explore this, we have calculated the lipid enrichment/depletion around the proteins in the inner leaflet of the ReLPS outer membrane using the methods described by Corradi et al [10]. Overall, in line with our findings from the analyses presented already, we find each protein has its own characteristic pattern of lipids around it (Figure 6B). In general there is no overall enrichment of PE nor PG lipids around the Omps. The greatest variations are seen with cardiolipin; while overall there is depletion of cardiolipin around the Omps (within a 1.4 nm radius from the centre of geometry of the protein). On the basis of electrostatics alone, one might expect PG and cardiolipin to behave similarly around the proteins; from our results it is evident that this is not the case for the six Omps studied here. The D/E index provides a view of the lipid levels within a given radius around the protein rather than details of specific regions within that radius. To get a more detailed characterisation of specific regions of lipid depletion or enrichment, contact maps of PG with each protein were determined for the 25-30 |as interval of each system (Figures 6A and S9). We then cross-referenced to determine persistent binding site between repeat simulations of the same protein. We found that in particular FhuA and BtuB had prominent PG binding sites. The three persistent bindings sites of FhuA mainly consisted of charge or aromatic residues. Interestingly, one of these binding sites is located directly below the LPS binding site resolved in the X-ray structure (PDB code = 1QFG)[17]. Similarly two persistent PG binding sites were observed in the lower leaflet of BtuB. For both BtuB and FhuA there were also additional patches of PG enrichment, but these were not consistent between repeat simulations in terms of size and location. Interestingly clustering of cardiolipins is seen beyond the 1.4 nm radius used to calculate the D/E index in some of the simulations. This effect is seen in a number of simulations and appears to be independent of protein type. Perhaps the most pronounced effect is seen in one of the simulations of FhuA (Figure 6); while cardiolipin is depleted within 1.4 nm of the protein, molecules of cardiolipin are clustered together in one region of the membrane beyond this distance. This cluster of cardiolipin molecules in the lower leaflet corresponds to a region of LPS depletion above it, in the upper leaflet. This compensation effect is also seen in simulations of OmpX and BtuB. We previously reported this phenomenon in simulations of OmpA and OmpF. Extension to four additional Omps in the present study reveals that the effect is common, but likely not specific. Given this effect is not seen in both independent repeats of the Omp simulations, other than FhuA, and it occurs outside of the annular lipids of the protein, this suggests that the compensation for LPS depletion, by cardiolipin clustering is a general feature of the outer membrane and is not dependent upon the protein type.

**Figure 6.**
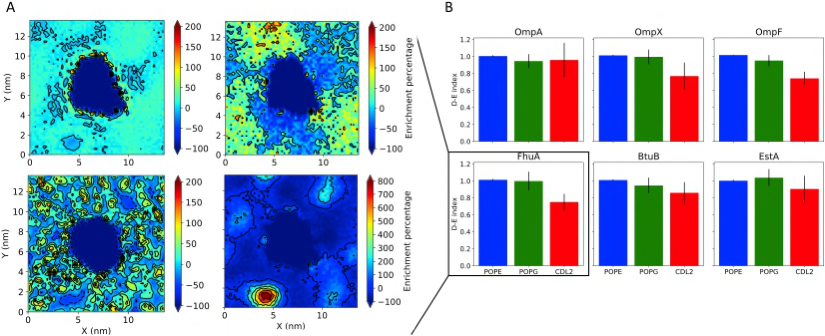
A) 2D lipid density analysis for the ReLPS outer membrane systems. Density maps were done for, bottom left) lipopolysaccharide (LPS), top left) 16:0-18:1 phosphoethanolamine (POPE), top right) phosphatidylglycerol (POPG) and bottom right) cardiolipin (CDL2) lipids and were averaged over 25-30 ps. The density maps are coloured by enrichment (>0) or depletion (<0) of each lipid type with respect to the average density of a given lipid type. B) Depletion-Enrichment (D-E) indices for POPE, POPG and CDL2 for each protein in the ReLPS membranes. The D-E index was obtained by dividing the lipid composition the 1.4 nm shell around the protein by the bulk membrane composition; therefore a D-E index > 1 indicates enrichment, while a D-E index < 1 indicates depletion. The D-E index was calculated from 10-30 ps in 5 ps blocks for both repeats to get eight values for each system; these eight values were then averaged and the error determined from their standard deviation.

### O-antigen orientation within the outer leaflet of the outer membrane

The O-Antigen chains attached to the RaLPS outer core act as a chemical and physical barrier to the permeation of molecule across the outer membrane. In previous sections of this paper, we have shown that the pattern of contacts of the six Omps with sugar moieties are protein dependant, here we ask if the Omp-sugar interactions impact upon the behaviour of the O-Antigen chain. To this end we calculated the end-to-end tilt of each O-Antigen chain with respect to the z-axis and distance from the protein. The distances are divided into ‘local’ and ‘bulk’, where local is defined as the lipid A section of each smooth LPS molecule being within 2.5 nm of the protein surface, as this is the approximate radius of the annular regions around most membrane proteins, see for example [10, 20]. The bulk region is defined as the region beyond the local region. The relative orientation of O-Antigen chains was determined by the calculation of the angle between end-to-end vectors of neighbouring O-Antigen chains. The angles took positive values if the chains tilt toward each other and negative values if they did not. In both cases the results were presented as probability distributions with respect to the angles. For the O-Antigen systems, bulk region sugar chains rarely have tilts angles greater than 45°, whereas in the local region, O-Antigen tilt angles can be as large as 65° (Figures 7 and S10A). Tilting of O-Antigen chains with respect to the z axis in the local region in general has a positive correlation with the size of the Omps; this observation was particularly pronounced for the OmpA and OmpF, respectively, where the O-antigen chains in the vicinity of OmpF had a considerably larger tilt angle than the chains nearer to the smaller protein, OmpA (Figure 7). It is interesting to note that EstA also showed a greater propensity of O-Antigen tilt at short-ranges even though the transmembrane region of EstA is not much larger than that of OmpA; this observation reflects the extensive interactions of the large extracellular region of EstA with the O-Antigen chains. The pairwise angles were all distributed from-50° to 50° with the most probable relative tilt being ~± 25°. Once the most probable angle is reached the distributions undergo a gradual decay to 0, highlighting the packed nature of these system. For every Omp, pairs of neighbouring O-Antigens are more likely to tilt towards each other (tenting), than they are away from each other; this leads to O-Antigen chains clustering together. Similarly, shorter, atomistic simulations reported by Im and co-workers shown tenting of O-Antigen chains around OmpF[21].

**Figure 7.**
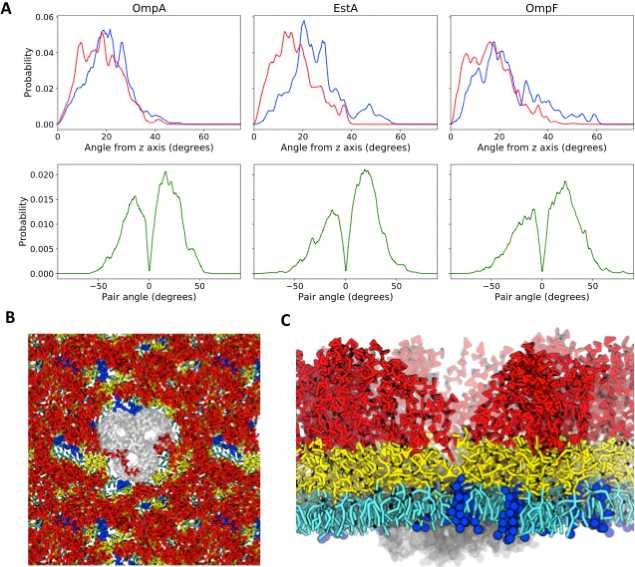
A) Probability distributions of (top) O-Antigen chain tilt with respect to the z axis and tilt between neighbouring pairs of O-Antigen chains for 25-30 ps of the OANT systems. The tilt of a single O-Antigen chain was calculated using its end to end vector. For the chain tilt with respect to the z axis, two probability distributions were generated for smooth lipopolysaccharides (LPS) closer than 2.5 nm (blue line) or further than 2.5 nm away from a given protein’s surface (red line). For the pairwise O-Antigen distributions if the chains tilted towards each other the angle was positive; in all other cases the angles would be negative. Snapshot of OANT_PE system with an embedded OmpF taken at 30 ps from C) above and D) the side of the membrane. Colour key: grey = protein red = O-Antigen chain, yellow = LPS core oligosaccharides, lipid A = cyan and gold and 16:0-18:1 phosphoethanolamine (POPE) lipids in the upper leaflet = blue.

The addition of POPE to the upper leaflet of the outer membrane results in an overall rise in the probability of tilt angles above 45°, especially within the local region of the protein (Figure S10B). Whereas the pair angles seem largely unchanged from the OANT membrane simulations, which maybe suggest that extreme tilts are mirrored by their local environment. Here we focus on the OmpF OANT_PE simulations, as these systems exhibited the largest differences in O-Antigen tilting behaviour when POPE was added to the outer leaflet. Snapshots taken at the 30 ps point from one of the simulations show a few O-antigen chains curved over the extracellular loops of OmpF, but only partially obstructing one of the pores of the trimer (Figure 7); an effect that was seen throughout the trajectory. We determined the average O-antigen contacts per frame with all residue numbers within each repeating unit (1-340) of OmpF of the system in Figure 5 from 25-30 ps (Figure S11). As in the atomistic study of OmpF in an O-antigen containing model the OM reported by Im and co-workers, we observed significant contacts with parts of the loops L4 (residues 160-172), L5 (residues 197-209), L6 (residues 237-253) and L8 (residues 318-330). Compared to the study of Im and co-workers, we see less extreme tenting of O-Antigen chains near OmpF; which is a likely consequence of the O-Antigen chains in this study being constructed of 4 monomer sugar units compared to 5 units in the work of Im, and thus it would not be unreasonable to observe increasing O-Antigen flexibility with an increase in chain length. In simulations of the other Omps presented here we did not observe O-antigen chains to block access to the pore of the Omps and rarely did they tilt over the protein. Furthermore, in simulations in which POPE was present in the outer leaflet, we observed O-antigen chains tilt away from small clusters or even single POPE lipids to preferentially interact with neighbouring sugars. Thus, despite the presence of a thick O-antigen layer in the outer leaflet of the outer membrane, on the basis of our simulations, this layer does not impede access to the proteins embedded within the membrane.

## Discussion

We have shown that the six outer membrane proteins we have studied here, OmpA, OmpX, FhuA, BtuB, OmpF and EstA have unique patterns of interaction with the outer membrane. In the inner leaflet we find that cardiolipin is depleted within the vicinity of all six Omps, in contrast to findings by ourselves and others that cardiolipin is enriched near some bacterial inner membrane proteins. We observe protein-independent clustering of cardiolipin molecules in the inner leaflet, directly beneath regions of lipopolysaccharide enrichment in the upper leaflet. We previously reported such behaviour close to OmpA and OmpF, the present study suggests this phenomenon is perhaps a more general characteristic of the outer membrane. In the outer leaflet we do not see any trends in protein interactions with LPS even in proteins that are similar in size or functionality. The Omp-LPS interactions vary across proteins, but also for a single protein they vary as a function of the level of lipopolysaccharide in the outer leaflet. For example interactions with basic residues dominate at the ReLPS level for most Omps, but at the RaLPS, in OmpX there are more interactions with acidic residues. These findings can in part be rationalised by the variations in the distribution of amino acids on the outer surface of these proteins. For example the extracellular loops of OmpA contain fewer lysine residues that point out towards the LPS than OmpX, despite the similar size of the proteins; both beta barrels contain just eight antiparallel strands. These compositional differences impact upon the orientations of the proteins within the different membrane environments as they dictate the pattern of interactions with LPS molecules. In the past it had been thought that the aromatics residues that are in general present in two bands, one near each mouth of the barrel of Omps anchor the protein into the headgroups of the lipids and then the barrels tilts to match the hydrophobic regions on their surface to the hydrophobic width of the surrounding membrane. We show that this is rather too simplistic a view, as the large extracellular loops and their interactions with the sugar moieties of the LPS molecules also play a role in Omp orientation. Furthermore the very different dynamics of the lipids in the two leaflets also impact upon the tilt angles of these proteins within the outer membranes. In the context of the in vivo outer membrane, which is known to be crowded by proteins, it is likely that the orientations of the proteins will also vary as a function of which other proteins are nearby, for the reasons already discussed; namely compositional differences. Interestingly we find that O-antigen chains preferentially tilt towards each other compared to towards Omps. We note that this may be important for recognition of Omps by other molecular species as they navigate through the ‘hairy’ layer of O-antigen chains of the outer membrane. It is important to place the current work in the appropriate biological context and discuss any limitations. For this study these are associated with protein sample size, and almost inevitably for molecular dynamics simulation, with simulation time and length scales. Of course there are very many more outer membrane proteins in *E. coli* than those studied here, thus it is possible that there may be some that are very similar in terms of their patterns of LPS interactions. The proteins we have chosen do cover a range of sizes and functions and while we are confident that the hypothesis of unique interactions will hold based on our findings here, it would be interesting to see similar studies of other Omps in future and indeed of Omps in other bacterial species too. The timescales we have achieved here (3040 gs per simulation) are state-of-the-art, and as the data on tilt angles reveals, they are absolutely necessary to achieve convergence for the slow-moving LPS molecules. Our simulations show some interesting features of the lipidic components of the outer membrane, namely the clustering of cardiolipins above regions of low LPS density in the outer leaflet and also the presence of clusters of PG lipids which appear to be independent of the protein they surround. In summary, the present study reveals that the six proteins studied here, have unique lipid fingerprints particularly with respect to lipopolysaccharide, and are not in general occluded or engulfed by O-antigen chains of the surrounding membranes.

## Experimental Procedures

### O-antigen parameterisation

The GROMOS 53A6 force field was used to perform simulations of duration 100 ns of *E. coli* O42 O-antigen chain units (with terminal glucose sugar), solvated with SPC water[22, 23]. Pressure was maintained at 1.0 bar using the Parrinello-Rahman barostat[24], while the temperature was held constant at 313 K using the Nose-Hoover thermostat[25, 26]. Electrostatic interactions were treated with the smooth particle mesh Ewald algorithm with a short-range cut-off of 0.9 nm, while the van der Waals interactions were truncated at 1.4 nm with long-range dispersion corrections for energy and pressure[27]. All bonds were constrained with the LINCS algorithm, enabling a timestep of 2 fs[28].

Subsequently, the united-atom particles were mapped into coarse-grain pseudoatoms, and equilibrium bond lengths and bond angles (for the coarse-grain O-antigen model) were determined from the relative conformations of these pseudoatoms, while the corresponding force constants were resolved from their dynamic movement. The coarse-grain system was then simulated in water for 100 ns and the resulting simulation data was compared with the comparable united atom simulations, enabling further fine-tuning of the coarse-grain parameter set. Thereafter the coarse-grain parameters (bond lengths, angles and dihedrals) were iteratively optimized as additional coarse-grain solvation simulations were performed with adjusted parameter sets until agreement was obtained between the similar coarse-grain and fine-grain solvation simulations within the range used for our original coarse grained model of LPS (additional details are provided in the supplementary information, Figure S12). The resulting O-antigen model and parameter set were combined with our pre-existing coarse-grain model and parameter set for *E. coli* RaLPS[29].

#### Membrane construction

Five different membrane models were simulated: 4 outer membrane systems and a 16:0 dipalmitoylphosphocholine (DPPC) system. The inner leaflet in all four of the outer membrane models was composed of 90% 16:018:1 palmitoylphosphoethanolamine (POPE), 5% 16:0-18:1 palmitoylphosphatidylglycerol (POPG) and 5% cardiolipin. The outer leaflet was comprised of either 100 % rough lipopolysaccharide (LPS) molecules; at either Re or Ra levels or smooth LPS which incorporates O-antigen, or a mixture of smooth LPS and POPE in a 4:1 ratio.

#### Simulation setup

Initial protein structures were obtained from the RCSB protein databank, they are OmpA: 1QJP[30], OmpX: 1QJ8[31], OmpF: 3POX[32], BtuB: 1NQE[33], FhuA: 1QFG [34] and EstA: 3KVN [35]. Any missing residues or broken loops where added with Modeller[36] and then the protein was given a transmembrane orientation determined by MemEmbed[37]. All initial membrane protein structures were generated with the Martini-maker module of CHARMM-GUI [38]. Each protein model was coarse-grained to the ElNeDyn model with an elastic network cut-off of of 0.9 nm and force constant of 500 kJmol^−1^nm^−2^ [39]. For each system the box height was set such that the distance above and below the membrane was 5 nm. For the smaller proteins the initial box dimensions in the xy plane were around 13×13 nm, while for larger proteins (OmpF and EstA) initial box dimensions were as large as 17×17 nm (see Table 1). The core of LPS was neutralised with Ca^2+^ ions, while the rest of the charge deficit was removed via the addition of Na+ ions. Following this 0.15 M of NaCl was added to each system to give biologically relevant salt concentrations.

#### Simulation protocols

All simulations were performed using the GROMACS package version 2016 and the Martini version 2.2 force field[40] [41]. All > simulations were carried out at 323 K with a stochastic velocity rescale thermostat with a coupling constant of 1.0 ps. Initial minimisations followed the CHARMM-GUI protocol[42], which involved an initial steepest decent minimisation followed by a series of equilibrations with 5 fs to 10 fs timesteps for a total of 20 ns with a Berendsen barostat (4.0 ps coupling constant)[43]. During these equilibrations the position of the protein was restrained with 1000 kJmol^−1^nm’^2^ harmonic restraints on the backbone atoms. Following this production runs of up to 40 ps (see Table 1) were generated with a 10 fs timestep and analysed as described elsewhere. For all production runs, the Parrinello-Rahman semi-isotropic barostat was used with a 12.0 ps coupling constant [24]. Repeat simulations were performed for all systems by assigning random velocities from the stage after which the initial system had been generated from CHARMM-GUI[38].

The Lennard-Jones potential was cut off using a Potential shift Verlet scheme at long ranges. The reaction field method was used for the electrostatics, with dielectric constants of 15 and infinity for charge screening in the short and long range regimes, respectively (as recommended for use with MARTINI2). The short range cut-off for both the non-bonded and electrostatic interactions was 1.2 nm. The non-bonded Neighbour lists were updated every 10 steps.

#### Analysis

All analysis was carried over 25-30g.s, unless stated otherwise, using a combination of GROMACS tools [ref] and in-house scripts which utilised the MDAnalysis python module [44]. To calculate the tilt of each protein the principal axis of inertia of the transmembrane region of the protein backbone, or for OmpF the backbone entire timer, was determined. The protein tilt was then obtained by the angle between the previously obtained principal axis and the z axis. Contact histograms and maps between a single protein and a range of target atoms were determined with in-house scripts that used a cut-off of 0.6 nm. If multiple contacts were present between a protein residue and a single target atom only one contact was recorded. The membrane thickness was determined with an in-house script that utilised Voronoi diagrams to get the membrane thickness, in the same manner as the APL@Voro analysis tool [45]. The thickness was measured using only the phosphate beads in each leaflet. The depletion/enrichment indices were determined by first counting the number of lipids with a center of geometry within 1.4 nm and then comparing this number to the number expected in the bulk of the membrane using the procedure described by Corradi et al [10]. The enrichment maps were generated by first determining the 2D density map of the membrane using tools developed by Castillo et al[46]. Following this enrichment was determined using the procedure described by Corradi et al [10]. The D-E index was obtained by dividing the lipid composition the 1.4 nm shell around the protein by the bulk membrane composition; therefore a D-E index > 1 indicates enrichment, while a D-E index < 1 indicates depletion. The D-E index was calculated from 10-30 ps in 5 ps blocks for both repeats to get eight values for each system; these eight values were then averaged and the error determined from their standard deviation (as in [10]). Any images of systems seen here were generated with VMD[47].

The tilt of each O-Antigen chain was determined by first generating an end to end vector for each chain that spanned from the sugar connected to the LPS core to the end of the chain (n.b this is a normalised vector). The tilt angle of each chain was the angle between the end to end vector of the chain and the z axis. Each chain was then split into two groups based on whether the center of geometry Lipid A headgroup of each LPS was closer than 2.5 nm to the protein surface. The values in a single group were then used to generate a probability distribution that was normalised to have a total integral of one. The pairwise O-Antigen angle probability distributions were determined by first finding all pairs of O-Antigen chains within a cut-off of 0.6 nm from each other. Then the angle between the end to end vectors of each pair of chains was determined. If the O-Antigen chains were tilting away from each other, the tilt angle was made negative; otherwise the tilt angle was positive.

